# Identification and molecular characterization of novel viruses in Ugandan cattle

**DOI:** 10.1101/2022.01.05.475030

**Authors:** Stephen Balinandi, Juliette Hayer, Harindranath Cholleti, Michelle Wille, Julius J. Lutwama, Maja Malmberg, Lawrence Mugisha

## Abstract

The risk for the emergence of novel viral zoonotic diseases in animals and humans in Uganda is high given its geographical location with high biodiversity. We aimed to identify and characterize viruses in 175 blood samples from cattle selected in Uganda using molecular approaches. We identified 8 viral species belonging to 4 families (*Flaviviridae, Peribunyaviridae, Reoviridae* and *Rhabdoviridae*) and 6 genera (*Hepacivirus, Pestivirus, Orthobunyavirus, Coltivirus, Dinovernavirus* and *Ephemerovirus*). Four viruses were novel and tetantively named as Zikole virus (Family: *Flaviviridae*), Zeboroti virus (Family: *Reoviridae*), Zebtine virus (Family: *Rhabdoviridae*) and Kokolu virus (Family: *Rhabdoviridae*). In addition, Bovine hepacivirus, Obodhiang virus, *Aedes pseudoscutellaris reovirus* and Schmallenberg virus were identified for the first time in Ugandan cattle. We report a broad range of viruses including novel ones in the blood of cattle likely as reservoir hosts for emergence of novel viruses with serious public health implications.

**Highlights:**

- Pan PCR and High Throughput Sequencing Approaches reveal novel viruses in the blood of cattle.
- Identified 8 viral species belonging to 4 families: *Flaviviridae, Peribunyaviridae, Reoviridae* and *Rhabdoviridae*
- 60% of the identified viruses were found in the Ankole cattle breed

## Introduction

Since the mid-1900s, infectious diseases of livestock have been identified, not only as a major threat to animal health, welfare and production, but also to the overall global health security especially in developing countries (Cleaveland et al., 2017; Grace D et al., 2012). Moreover, more than 60% of documented emerging, and re-emerging human diseases are zoonotic in nature. That is, viruses that are endemic within animal populations spill over into humans through vectors and other transmission pathways and cause epidemics (Morse, 2004), or global pandemics as with SARS-CoV-2 (R. Lu et al., 2020)

Uganda is a known hotspot for the emergence of novel pathogens due to its conducive climatic conditions and high biodiversity that include a variety of wild and domestic animals, birds and insects acting as disease vectors, hosts or reservoirs, and the attendant anthropogenic factors such as deforestation (Allen et al., 2017; Jones et al., 2008). Indeed, in just over 10 years, several new high consequence viral pathogens have been identified, including *Bundibugyo ebolavirus* (Family: *Filoviridae*) (Towner et al., 2008), *Sosuga pararubulavirus* (Family: *Paramyxoviridae*) (Albariño et al., 2014), Bukakata orbivirus (*Chobar George virus*; Family: *Reoviridae*) (Fagre et al., 2019) and Ntwetwe virus (Family: *Peribunyaviridae*) (Edridge et al., 2019). Other viruses of public health importance that were previously discovered in Uganda, especially during the middle of the 20^th^ century, include *Zika virus*, *West Nile virus* and *O’nyong-nyong virus* (Dick, Kitchen, & Haddow, 1952; Smithburn, Hughes, Burke, & Paul, 1940; Williams, Woodall, & Gillett, 1965).

Despite enhanced efforts to identify and manage potential disease-causing pathogens in Uganda through improved surveillance, increased public and health sector awareness, and expedited transportation of suspected samples to the testing laboratories (Borchert et al., 2014; Kemunto et al., 2018; Kiyaga et al., 2013; Shoemaker et al., 2018), the disease burden associated with undifferentiated illnesses of unknown origin remains high (Lamorde et al., 2018). As such, it is critical to implement broad investigations of potential disease-causing agents to include animal hosts and vectors to identify the potential sources of viral pathogens likely to cause diseases in both animals and humans. These investigations would align well with the WHO’s Priority ‘Disease X’, which represents a currently unknown serious global health threat that can emerge in humans (Mehand, Al-Shorbaji, Millett, & Murgue, 2018). In our current study, we aimed at identifying and characterizing viruses from blood samples collected in cattle from selected districts of Uganda. Previous studies have alluded that bovines are a major source of zoonoses (McDaniel, Cardwell, Moeller, & Gray, 2014). More specifically, cattle have been implicated as hosts for 2 human viruses. Firstly, the dated emergence of Human coronavirus OC-43 in 1890 coincides with a pandemic of human respiratory disease in humans between 1889 – 1890, and it is believed that this was after the virus emerged following a cross species transmission event of Bovine coronavirus (Crookshank, 1897; Vijgen et al., 2005). Secondly, measles virus is said to have crossed to humans after it diverged from rinderpest virus, a historical bovine virus, around the 11^th^ to 12^th^ Century (Furuse, Suzuki, & Oshitani, 2010). Therefore, with the high consumption of meat and other dairy products, as the main source of food in Uganda (UBOS, 2017), it is prudent to understand the viral diversity in this animal reservoir to expand on our pandemic preparedness toolkit.

## Materials and Methods

### Ethics Statement

The authors confirm that the ethical policies of this journal, as noted on the author guidelines page, have been adhered to: sample collection was approved by the Institutional Animal Care and Use Committee (IACUC), School of Veterinary Medicine & Animal Resources (SVAR), Makerere University (Reference Number: SVARREC/03/2017) and the Uganda National Council for Science and Technology (UNCST) (Reference Number: UNCST A580). In addition, written consent was obtained from all animal owners, or their representative(s), following a detailed explanation of the study objectives.

### Study Sites

Uganda is an East African country located along the Equator, lying between 4°N and 1°S latitudes, and between 30°E and 35°E longitudes. Thus, many areas within the country have a typical tropical climate; experiencing relatively humid conditions (average of 70%), rainfall (700-1,500 mm) and daily temperatures that are between 20-30°C throughout the year (UBOS, 2017). Its natural vegetation is varied although it is mainly open wooded savannah grassland in most of the country, thus making livestock keeping a major agro-economic activity for the country. Recent data from the Ministry of Agriculture, Animal Industry and Fisheries (MAAIF) and Uganda Bureau of Statistics (UBOS), show that Uganda has over 14 million cattle, 16 million goats, 5 million sheep and 4 million pigs with every household owning at least one type of livestock (UBOS, 2017).

This study was conducted using blood samples from cattle that were sampled from 5 purposively selected districts of Uganda: Kasese (0°N, 30°E), Hoima (1°N, 31°E), Gulu (3°N, 32°E), Soroti (1°N, 33°E) and Moroto (2°N, 34°E) (Figure 1 and Table 2). These districts were selected as they represent areas with distinct agroecological variations (Drichi & National Biomass, 2003; Wortmann & Eledu, 1999). Briefly, Kasese and Moroto districts lie at the extreme ends of the western and eastern borderlines, respectively, of the Ugandan Cattle Corridor – a large broad zone of semi-arid rangelands, where cattle-rearing is a preoccupation of the local communities. Gulu and Soroti districts lie within the swampy lowlands of the Lake Kyoga basin, while Hoima district is a medium altitude area with mixed agricultural practices.

**Figure 1:**
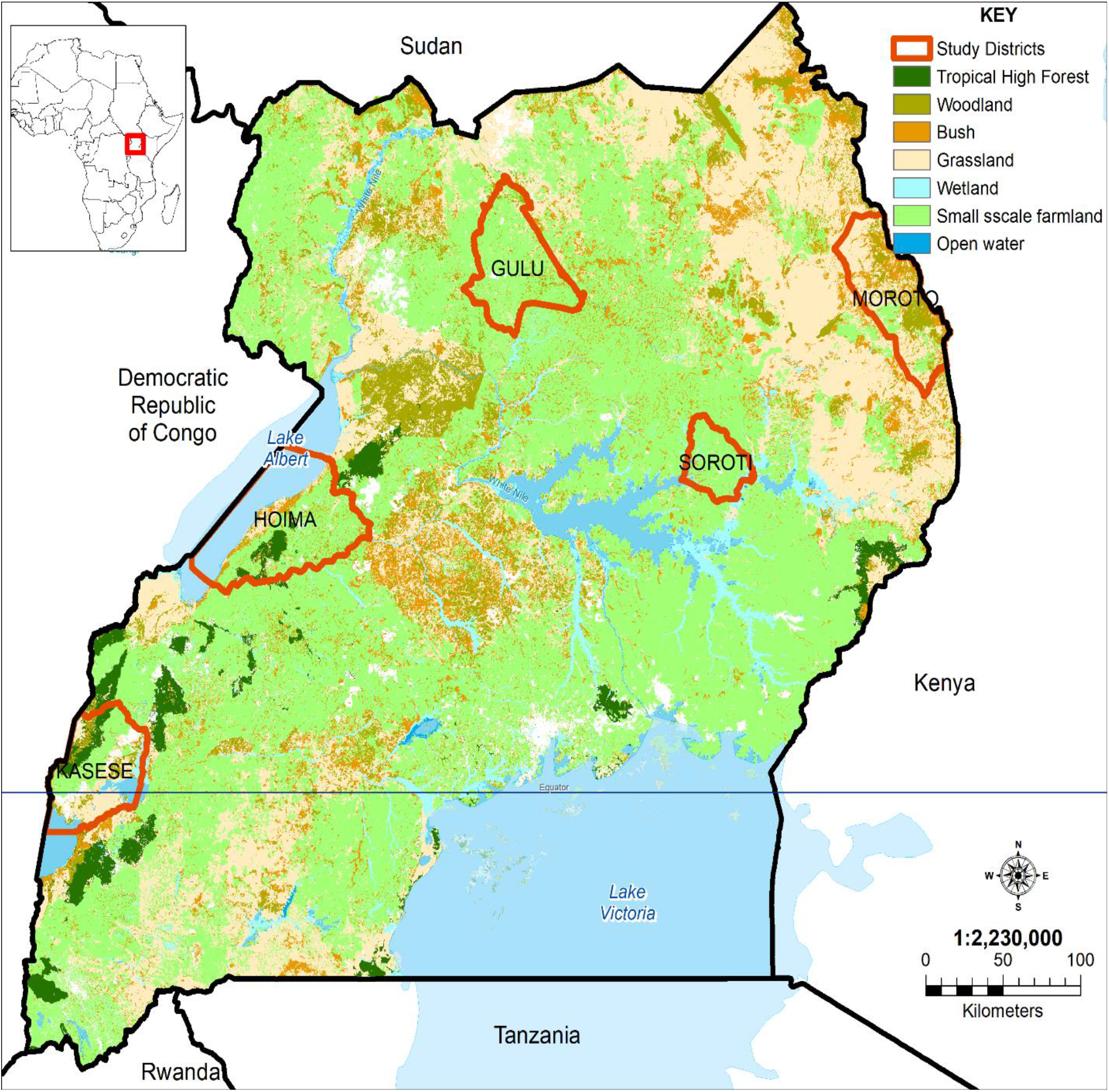
Map of Uganda showing the sampling sites in this study (Source: This map was created using open-source data in ArcGIS software, v10.2, Environmental Systems Research Institute, Inc., Redlands, CA, USA). Inset is a map of Africa with the enlarged region boxed in red.

### Study Samples

In this study, we used samples collected as part of a multidisciplinary baseline investigation on tickborne disease in cattle in Uganda (Balinandi et al., 2021; Malmberg & Hayer, 2019). A subset of 175 cattle samples was randomly selected from a total of 500 sampled animals, comprising 35 samples from each of the 5 study districts (Gulu, Soroti, Moroto, Hoima, and Kasese). In the field, information about each of the sampled animal such as their age, sex, breed, health status, tick burden, grazing system, etc., was collected at the time of sample collection using a standardized biodata form. In addition, about 4ml of whole blood was drawn from the jugular or tail veins of restrained cattle into EDTA tubes (BD Vacutainer®, Franklin Lakes, NJ, USA). Blood was then immediately separated into small aliquots of whole blood and serum, which were immediately frozen in liquid Nitrogen before they were transported to Uganda Virus Research Institute (UVRI), Entebbe, Uganda, and stored at −80°C until RNA extraction and family-wide conventional pan-PCR was undertaken.

### pan-PCR

Total nucleic acids were extracted from whole blood samples using the MagMax™ 96 viral Isolation kit (Applied Biosystems, Vilnius, Lithuania), according to the manufacturer’s instructions. Extracted RNA was transcribed into cDNA using Superscript III First-strand Synthesis Supermix (Invitrogen, CA, USA). We assayed the samples for 5 viral taxa: *Filoviridae*, *Flaviviridae*, *Alphavirus*, *Rhabdoviridae*, and *Orthobunyavirales*, using protocols previously developed and applied for the USAID/PREDICT II program (Moureau et al., 2007; PREDICT, 2014; Sanchez-Seco, Rosario, Quiroz, Guzman, & Tenorio, 2001; Zhai et al., 2007) (Table 1). These viral taxa also represent the main groups of viruses from which most members of arboviruses are frequently identified (Hubálek, Rudolf, & Nowotny, 2014; Woolhouse & Gowtage-Sequeria, 2005).

**Table 1.**
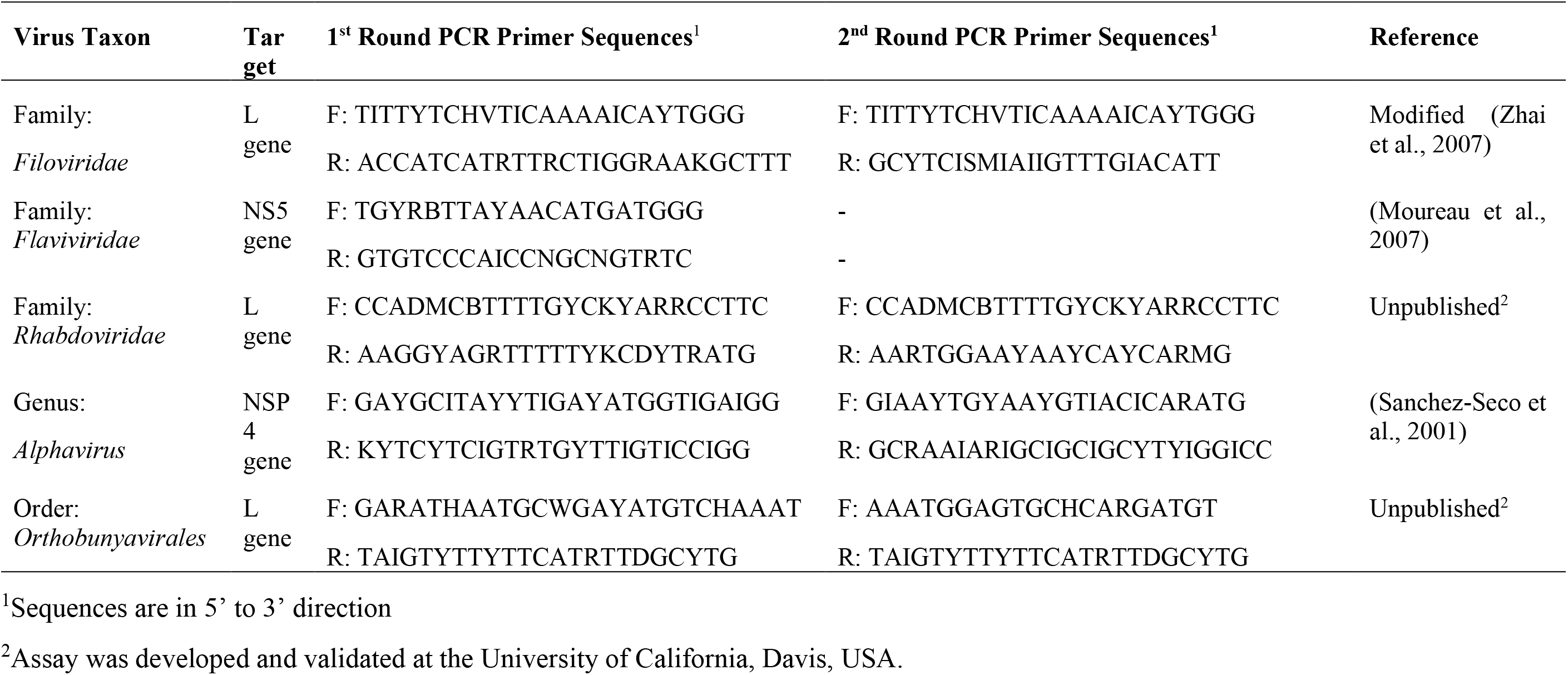
Pan-PCR Primer Sequences and Protocols used for Virus Detection in Cattle, Uganda, 2017.

### High-throughput sequencing

Ten serum samples (Table 2), corresponding to the whole blood samples that were positive for any viral family, including those that showed non-specific bands on gel electrophoresis, were preserved in TRIzol™ Reagent (Invitrogen, Thermo Scientific) and shipped on dry ice to the Swedish University of Agricultural Sciences (SLU), Uppsala, Sweden, for high-throughput sequencing.

**Table 2:** Characteristics of Pan-PCR Positive Cattle, Uganda, 2017.

| Animal ID# | Sample Collection Area |  | Animal Characteristics |  |  |  | Results of Pan-PCR |
| --- | --- | --- | --- | --- | --- | --- | --- |
|  | District | GPS Location | Age<sup>a</sup> | Sex | Breed | Health |  |
| G04B | Gulu | N 03.03243, E 032.30449 | 2 | M | Ankole | Good | Positive (Rhabdovirus) |
| G11A | Gulu | N 03.01411, E 032.28941 | 6 | F | Ankole | Good | Positive (Orthobunyavirus) |
| H11A | Hoima | N 01.64420, E 031.29474 | 5 | F | Ankole | Good | Indeterminate (Orthobunyavirus) |
| H26B | Hoima | N 01.32115, E 031.23425 | 2 | F | Zebu | Good | Indeterminate (Orthobunyavirus) |
| K09B | Kasese | S 00.10048, E 029.82413 | 8 | F | Ankole | Good | Indeterminate (Orthobunyavirus) |
| K16B | Kasese | S 00.04747, E 029.82864 | 2 | F | Ankole | Good | Indeterminate (Orthobunyavirus) |
| K28A | Kasese | N 00.29223, E 030.16044 | 1 | M | Ankole | Good | Positive (Orthobunyavirus) |
| M15B | Moroto | N 02.53587, E 034.53321 | 1 | F | Zebu | Good | Positive (Rhabdovirus) |
| S37B | Soroti | N 01.92280, E 033.49659 | 1 | M | Zebu | Good | Indeterminate (Orthobunyavirus) |
| S41B | Soroti | N 01.93135, E 033.49284 | 1 | M | Zebu | Good | Positive (Rhabdovirus) |

RNA was extracted using the GeneJet RNA purification kit (Thermo Scientific^TM^), as per the manufacturer’s instructions. The eluted serum RNA was further concentrated by RNeasy MinElute Cleanup Kit (Qiagen) before the RNA was randomly amplified with the Ovation RNA-Seq System v2 (Tecan) following the manufacturer’s recommended protocols. The concentration and size distribution of generated double-stranded cDNA products were estimated by the TapeStation 4200 system (Agilent Technologies). Sequencing libraries were prepared from 50ng of DNA according to the manufacturer’s preparation guide #112219, using the SMARTer ThrupPLEX DNA-Seq kit (R400676, Takara-Clontech). Briefly, DNA was fragmented using a Covairs E220 system, aiming at 350-400bp fragments. The ends of the fragments were end-repaired, and stem-loop adapters were ligated to the 5’ ends of the fragments. The 3’ end of the stem-loop was subsequently extended to close the nick. Finally, the fragments were amplified, and unique index sequences were introduced using 8 cycles of PCR, followed by purification using AMPure XP beads (Beckman Coulter). The quality of the library was evaluated using the Agilent 2200 TapeStation system and a D1000 ScreenTape-kit. The adapter-ligated fragments were quantified by qPCR using the Library quantification kit for Illumina (KAPA Biosystems/Roche) on a CFX384Touch instrument (BioRad), prior to cluster generation and sequencing.

Thereafter, a 300 pM pool of the sequencing libraries in equimolar ratio was subjected to cluster generation and paired-end sequencing, with 150bp read length in an SP flow cell and the NovaSeq6000 system (Illumina Inc.) using the v1 chemistry, according to the manufacturer’s protocols. Base-calling was done on the instrument using RTA 3.3.4 and the resulting .bcl files were demultiplexed and converted to FASTQ format with tools provided by Illumina Inc., allowing for one mismatch in the index sequence. Additional statistics on sequence quality were compiled with an in-house script from the Fastq-files, RTA and CASAVA output files. Sequencing was performed by the SNP&SEQ Technology Platform, part of the SciLifeLab, in Uppsala, Sweden. Sequence libraries have been deposited in SRA under the study PRJEB46076 with Accession Numbers: ERR6496566, ERR6496567, ERR6496875, ERR6496876, ERR6496901, ERR6496964, ERR6497048, ERR6497050, ERR6497350 and ERR6497480.

### Bioinformatics

Quality control and trimming of the raw sequencing reads was performed using FASTP (Lipman & Pearson, 1985). The good quality reads were then aligned to the cattle genome (Bos_taurus.ARS-UCD1.2.; www.ncbi.nlm.nih.gov/assembly/GCF_002263795.1/) using Bowtie2 (Langmead & Salzberg, 2012) to eliminate reads from the host. The remaining reads were *de novo* assembled using MEGAHIT (Li et al., 2016). The obtained assembled contigs were taxonomically classified using two methods: KRAKEN 2 (Wood, Lu, & Langmead, 2019) and DIAMOND (Buchfink, Xie, & Huson, 2015), both using the NCBI non-redundant protein database *nr* (release of April 2019) (Deng et al., 2006). This analysis was also performed at reads level, on the reads remaining unmapped after the alignment to the cattle genome.

The longest assembled contigs (length>1000 nt) that were classified as viral were retrieved for further analyses. BLASTN and BLASTX (Altschul et al., 1997; Chen, Ye, Zhang, & Xu, 2015) were performed to confirm the closest homologues and to check the alignments with viral sequences from the database. For some identified viruses for which only partial genomes were obtained by de novo assembly, all reads from the dataset of interest were mapped on the reference genome of the closest virus found in the taxonomic classification step. Prokka (Seemann, 2014) was also used to predict and annotate open reading frames from these longest contigs. Viral sequences generated in this study have been desposited in GenBank with Accession numbers as shown in Table 3

**Table 3.** Viruses identified by Metagenomics from Pan-PCR Positive Cattle, Uganda, 2017.

| Animal ID | Contig Characteristics |  |  |  |  | Identified Virus |  |  |
| --- | --- | --- | --- | --- | --- | --- | --- | --- |
|  | No. | Length (bp) | Full Length? | BLASTN Pident (%) | BLASTX Pident (%) | Virus Family (Genus) | Virus Species (Name given from this study) | GenBank Accession Numbers |
| H11A | 3 | 1,140 - 3,019 | - | 85.9 | - | *Flaviviridae, (Hepacivirus)* | *Bovine hepacivirus* | CAJZAA010000001-CAJZAA010000003 |
| G04B | 1 | 8645 | - | 87.2 | - | *Flaviviridae, (Hepacivirus)* | *Bovine hepacivirus* | OU592967 |
| K16B | 2 | 5,421 - 6,972 | Yes | 74.5 | - | *Flaviviridae, (Pestivirus)* | Novel pestivirus (Zikole virus) | OU592965 |
| K28A | 4 | 1,218 - 3,074 | No | 91.1 | 97.6 | *Peribunyaviridae, (Orthobunyavirus)* | *Schmallenberg virus* | CAJZAJ010000001-CAJZAJ010000004 |
| S37B | 1 | 2273 | No | 84.5 | 94.4 | *Reoviridae, (Coltivirus)* | Novel Coltivirus (Zeboroti virus) | OU592974 |
| H26B | 2 | 1,031 - 1,410 | - | 99.2 | 100 | *Reoviridae, (Dinovernavirus)* | *Aedes pseudoscutellaris reovirus* | CAJZAK010000001-CAJZAK010000002 |
| K09B | 4 | 1,013 - 1,167 | - | 98.2 | - | *Reoviridae, (Dinovernavirus)* | *Aedes pseudoscutellaris reovirus* | CAJZAC010000001-CAJZAC010000004 |
| M15B | 1 | 14618 | Yes | 96.4 | 98.3 | *Rhabdoviridae, (Ephemerovirus)* | *Obodhiang virus* | OU592970 |
| G04B | 1 | 14630 | Yes | 78.3 | 90.5 | *Rhabdoviridae, (Ephemerovirus)* | Novel Ephemerovirus (Kokolu virus) | OU592964 |
| S41B | 3 | 1,400 - 12,917 | - | 78.0 | 86.7 | *Rhabdoviridae, (Ephemerovirus)* | Novel Ephemerovirus (Zebtine virus) | CAJZAB010000001-CAJZAB010000003 |

### Phylogenetic analysis

Reference sequences were downloaded from RefSeq (www.ncbi.nlm.nih.gov/refseq/) and in addition to the top blast hits, amino acid or nucleotide sequences, were aligned using MAFFT (Katoh & Standley, 2013) in Geneious Prime (www.geneious.com/), and gaps trimmed using trimAL (Capella-Gutiérrez, Silla-Martínez, & Gabaldón, 2009). Phylogenetic trees were constructed using IQ-TREE (Nguyen, Schmidt, von Haeseler, & Minh, 2015) with the best model of amino acid or nucleotide substitution.

Based on BLAST results and phylogenetics, viral contigs were not included in the analysis if they were likely contaminants. For example, detecting viruses in clades dominated by arthropod viruses were not included. Furthermore, short contigs that blasted only against highly conserved 3’ ends of viruses likely represented misassignment. Finally, we cross-referenced our data with well-known lists of viral contaminants (Asplund et al., 2019). For all virus taxa, trees were constructed using the ORF containing the RNA-dependant RNA polymerase (RdRp). For the flaviviruses, we also included NS3 non-structural polyprotein, as this previously has also been found suitable (Atif et al., 2016; Rice, 1996).

We considered viruses to be novel if they had a <90% RdRp protein identity, <80% genome identity (Shi et al., 2018; Shi et al., 2016), and/or following cut-off thresholds specified by the International Committee on the Taxonomy of Viruses (ICTV) for each viral family (Lefkowitz et al., 2018).

## Results

### pan-PCR screening of viruses in blood samples from Ugandan cattle

To understand the degree of viral diversity and presence of specific viruses we screened 175 blood samples obtained from cattle in Uganda using a pan-PCR approach. The cattle ranged in age from <1-12 years (Median, 3.5; Mean, 3.7), and the breeds were Zebu (64.7%), Ankole (27.7%) and others (7.6%). At the time of sample collection, sampled cattle had rectal temperatures ranging from 36.0 – 40.6°C, and an average of 68 ticks per animal (Range: 0 – 450). However, the cattle from which we detected viruses were mainly young ones with an average age of 2.9 years (Range = 1-8), 60% of which were of the Ankole breed. These cattle had a normal rectal temperature (Range: 37.3 – 38.2°C) and a low to moderate tick burden (Range: 2 – 132 ticks).

Using a pan PCR approach, we found that 2.9% (5/175) of the samples were positive for either of 2 viral taxa. These included orthobunyaviruses (n=2) from samples collected in Kasese and Gulu districts, and rhabdoviruses (n=3) from samples collected in Gulu, Soroti and Moroto districts. In addition, we detected non-specific (indeterminate) electrophoretic bands in 5 samples that were obtained from animals in the districts of Hoima (n=2), Kasese (n=2) and Soroti (n=1), particularly for orthobunyaviruses (Figure 2).

**Figure 2.**
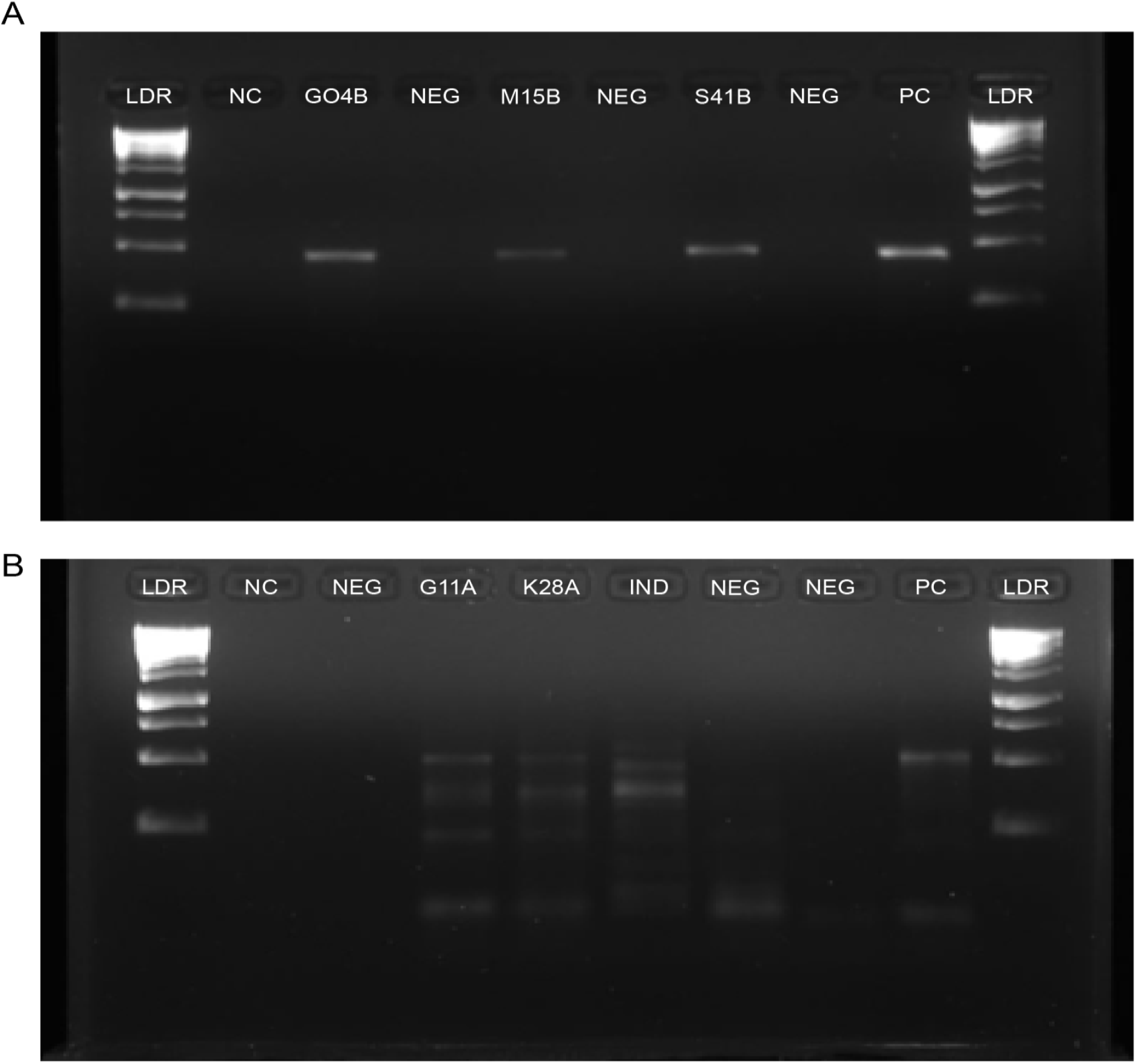
Gel images for some selected positive cattle samples (A) *Rhabdoviridae* and (B) *Orthobunyavirales*. Animal samples names correspond to G04B, M15B, S41B, G11A and K28A. LDR, NC, PC and IND comprise of the DNA Ladder, Negative Control, Positive Control, and the ‘Indeterminate’ pan-PCR samples, respectively.

### Large diversity of viruses revealed using metagenomics

To better understand the viral diversity, we performed metagenomic analysis on serum samples corresponding to the 10 positive pan-PCR whole blood samples. Of these 10 serum samples, we found evidence of at least one virus in 9 samples (Table 3). In one sample (G11A in Table 2) that had been collected from Gulu district, and was initially pan-PCR positive for orthobunyaviruses, a 382bp contig had a 98.9% nucleotide similarity to *Crimean-Congo hemorrhagic fever orthonairovirus*. Upon further analysis, these short contings also had high similarity to reverse genetic vectors (KJ648916, 98.91% identity, evalue 1e-85), and had low similarity and large gaps when aligned to other *Crimean-Congo hemorrhagic fever orthonairoviruses*, and hence this contig was removed from further data analysis. From the remaining 9 samples, metagenomic sequencing identified a total of 8 viruses, belonging to 4 families and 6 genera (Table 3). Importantly, this metagenomic approach confirmed the presence of virus positive samples generated in the pan-PCR approach.

Despite the pan-PCR approach failing to identify flaviviruses in the samples, metagenomics identified viruses belonging to this family in 3 of the submitted samples. These samples had showed non-specific gel bands on conventional pan-PCR. The identified flavivirus species were Bovine hepacivirus (Species *Hepacivirus N*, Genus *Hepacivirus*) in samples collected from Gulu and Hoima districts, and a novel flavivirus most closely related to Bat pestivirus (Genus *Pestivirus*) from Kasese district. This new flavivirus, which we have tentatively named Zikole virus, shares 74.5% nucleotide and 81% amino acid similarity to Bat pestivirus (Figure 3). Phylogenetically, Zikole virus is highly divergent to the clades containing pestiviruses from cattle, swine, and other ungulates (Figure 3).

**Figure 3.**
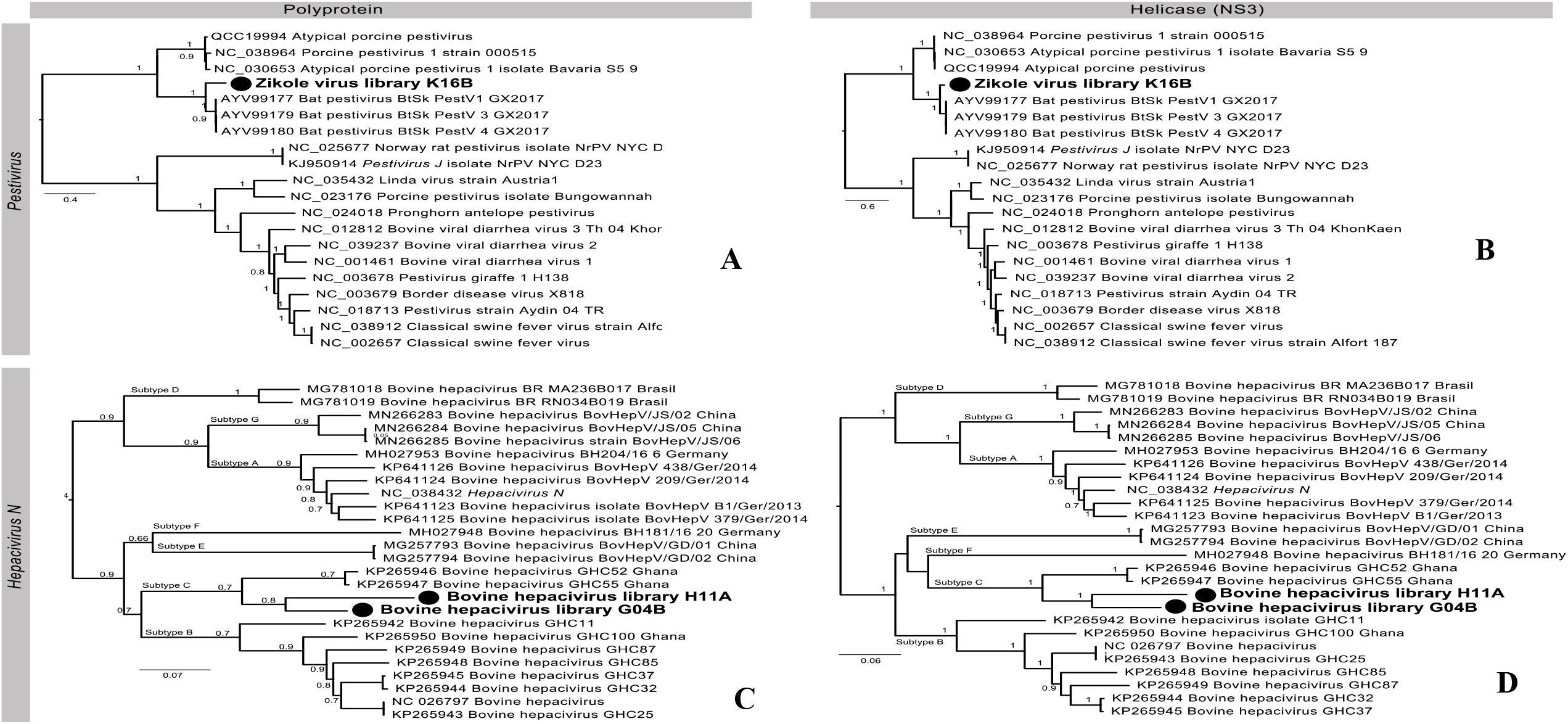
Phylogeny of select taxa of the *Flaviviridae.* Amino acid phylogeny (A, B) and nucleotide phylogeny (C, D) of *Pestivirus* (Zikole virus) and *Hepacivirus N* (Bovine Hepacivirus). Phylogenies in A, C are of the polyprotein and B, D are of the NS3 Helicase. For Pestivirus, the tree was midpoint rooted for clarity only, and for the Bovine Hepacivirus, the tree was rooted as per Lu *et al* (2019), including subtype information. Scale bar indicates number of substitutions per site. Sequences in this study are denoted by a filled circle.

Sequencing revealed three members of *Rhabdoviridae* belonging to the genus *Ephemerovirus*. These 3 viruses were Obodhiang virus (Species *Obodhiang ephemerovirus*) detected in a sample from Moroto district, and two novel viral species – one related to Puchong virus from a sample collected in Gulu district that we have tentatively named Kokolu virus, and another related to Kotonkan virus (Species *Kotonkan ephemerovirus*) in a sample from Soroti district that we have tentatively named Zebtine virus (Figure 4). The Kokolu virus shares 78.3% identity at the nucleotide level and 90.5% at the amino acid level across the L gene (containing the RdRp) with Puchong virus, while Zebtine virus shares 78.0% identity at the nucleotide level and 86.7% at the amino acid level, across the L gene with Kotonkan ephemerovirus. As per the current ICTV guidelines for species demarcation, rhabdovirus species must have a minimum of 15% amino acid sequence divergence in the L gene (ICTV, 2017). As such, the Zebtine virus identified in this study, falls well outside that threshold, but the status of Kokolu virus is less clear given its high nucleotide divergence, despite its relatively low (9.5%) amino acid divergence. The strain of Obodhiang virus identified in this study represents the first detection in Uganda since its description in mosquitos collected from South Sudan in 1963 (Schmidt, Williams, Lule, Mivule, & Mujomba, 1965), substantially expanding its geographic range.

**Figure 4.**
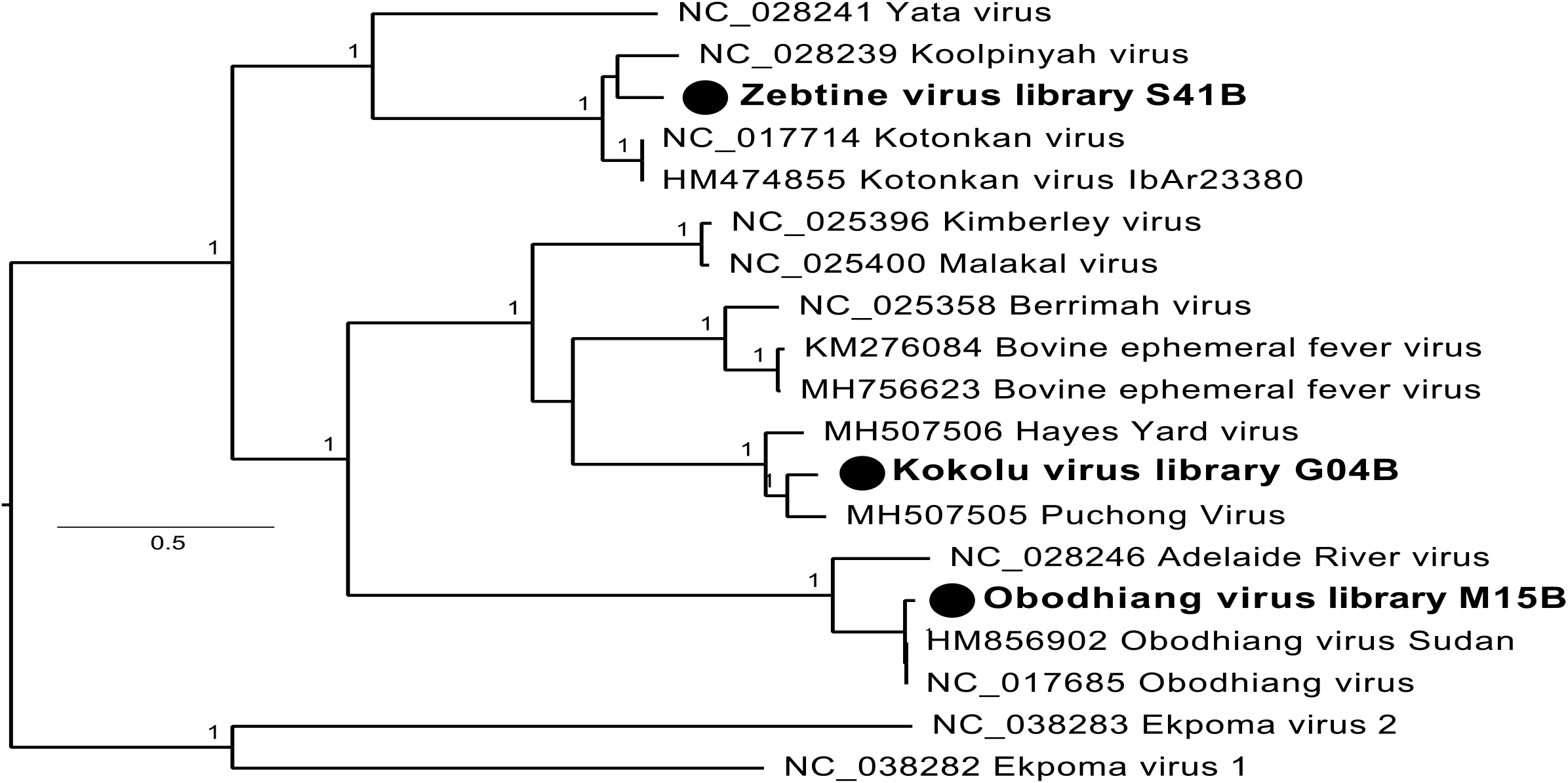
Phylogeny of the RdRp (L) protein of genus *Ephemerovirus* (Family: *Rhabdoviridae*). Ekoma virus 1 and 2 (genus *Tibrovirus*) are set as the outgroup. Scale bar indicates number of amino acid substitutions per site. Sequences in this study are denoted by a filled circle.

We also identified Schmallenberg virus (Species *Schmallenberg orthobunyavirus,* Genus *Orthobunyavirus,* Family *Peribunyaviridae*) from a sample collected from Kasese district, with a 91.1% nucleotide and 97.6% amino acid sequence identity in the RdRp gene (Figure 5). Across the L segment, the virus revealed here is sister to the clade containing Shamonda and Schmallenberg viruses, although the M segment falls into the clade comprising Shamonda virus, and more distantly related to Schmallenberg virus (Figure 5).

**Figure 5.**
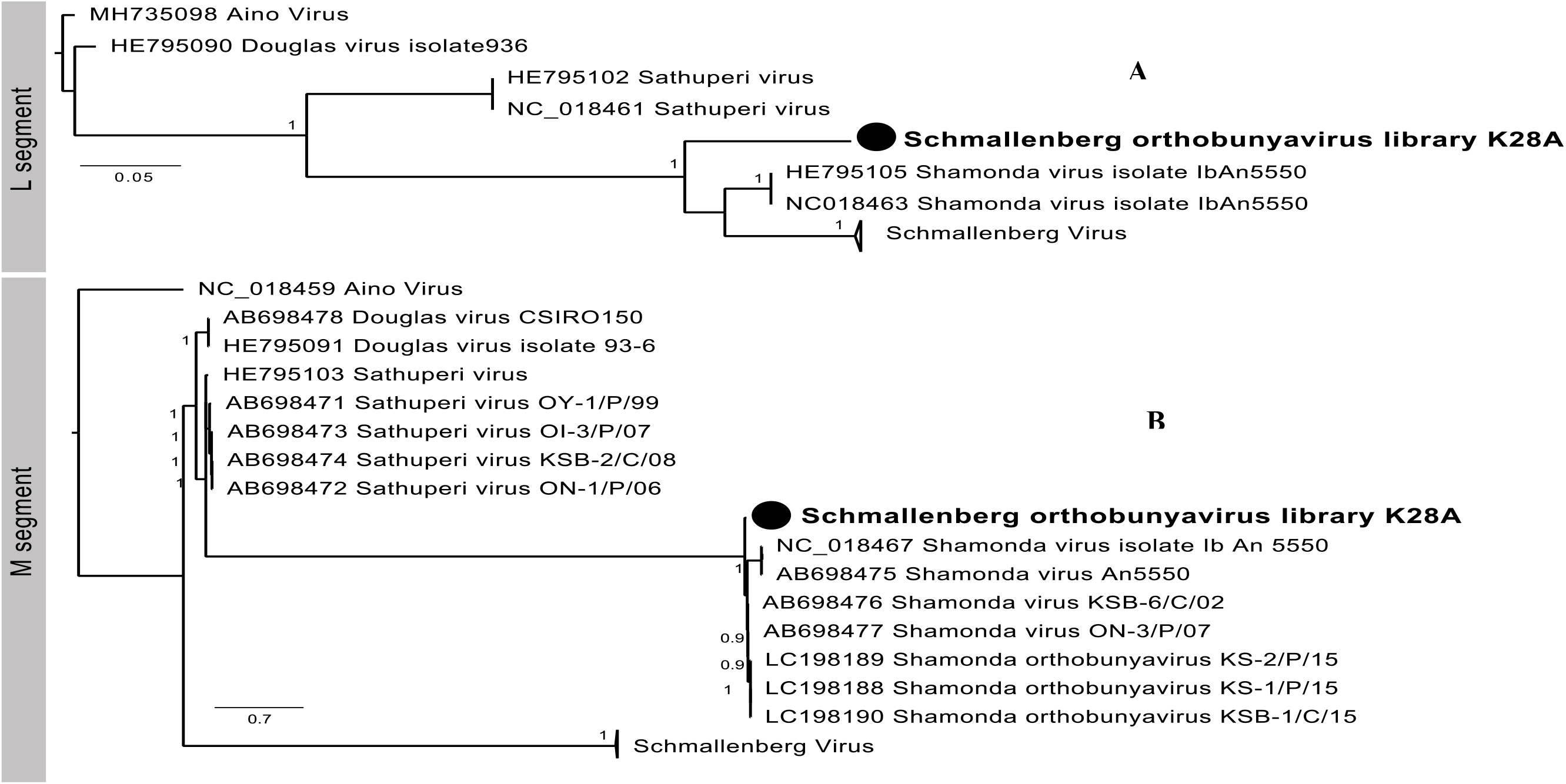
Phylogeny of the (A) L segment and (B) M segment of *Schmallenberg orthobunyavirus* including Schmallenberg Virus, Shamonda Virus, Douglas Virus and Sathuperi virus. Scale bar indicates number of nucleotide substitutions per site. Sequences in this study are denoted by a filled circle.

The final two virus species identified in this study belong to the Family *Reoviridae* (Figure 6); the first, which we have tentatively called Zeboroti virus, is a novel virus closely related to Tarumizu tick virus (Species *Tarumizu Coltivirus*, Genus *Coltivirus*) in a sample from Soroti district, with 84.5% nucleotide and 94.4% amino acid identity. According to the ICTV species demarcation guidelines for this genus, viruses belong to the same species if they have >89% nucleotide identity in segment 12, or amino acid identities of >55%, >57% and >60% in VP6, VP7 and VP12, respectively (Attoui et al., 2011). Unfortunately, we were only able to find genomic sequences belonging to segment 1 (out of the 12 putative segments), and therefore, ascertaining whether this virus is novel is challenging. Second, we identified Aedes pseudoscutellaris reovirus (Species *Aedes pseudoscutellaris reovirus*), the prototype-species for the genus *Dinovernavirus* of the *Reoviridae* family in two samples (one from Kasese district and another from Hoima district) with 98.2% and 100% nucleotide and amino acid identity, respectively. This virus is not well characterized, and it is unclear whether this virus is an arbovirus, an insect-specific virus, or a contaminant.

**Figure 6.**
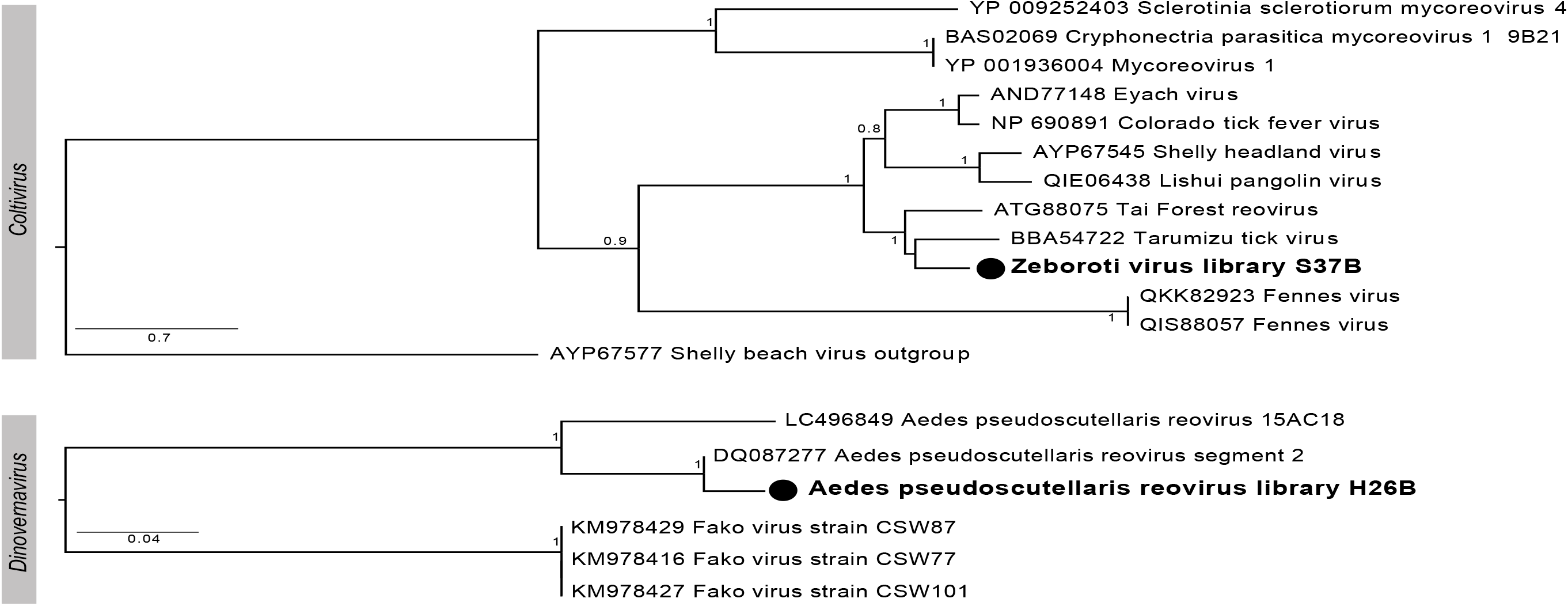
Phylogeny of *Reoviridae*. (A). Segment 1 of Coltivirus. Shelly beach virus was set as the outgroup. Scale bar indicates number of amino acid substitutions per site. (B) Segment 2 of Dinovernavirus (*Reoviridae*). The tree was mid-point rooted for clarity only. Scale bar indicates number of nucleotide substitutions per site. Sequences in this study are denoted by a filled circle.

## Discussion

This study investigated RNA viruses in blood samples that were collected from cattle in 5 districts in Uganda. Overall, we identified 8 viral species, belonging to 4 distinct viral families (*Flaviviridae, Rhabdoviridae, Peribunyaviridae* and *Reoviridae*). Four of the identified viral species in this study were novel (Zikole virus, Zeboroti virus, Kokolu virus and Zebtine virus), while the 4 other viruses (Bovine hepacivirus, Schmallenberg virus, Aedes pseudoscutellaris reovirus, Obodhiang virus), are being reported for the first time in Uganda. Thus, our findings, perhaps, are an early indication that cattle in Uganda may be harbouring more viruses than previously known, in line with similar high levels of viral diversity in cattle reported elsewhere (Dacheux et al., 2019; Kato et al., 2016; Kwok, Nieuwenhuijse, Phan, & Koopmans, 2020; McDaniel et al., 2014).

Interestingly, a substantial proportion of the identified and described viruses in this study are transmitted by, or fall into clades, associated with arthropods. Specifically, the coltivirus (Zeboroti virus) described here is most closely related, and fall within a clade dominated by tick-borne viruses, some of which are found in many parts of North America and Europe, where they cause acute febrile human illnesses (Fujita et al., 2017; Geissler et al., 2014; Hubálek & Rudolf, 2012). Other viruses detected in this study were mainly those currently associated with mosquito and culicoid flies (Yanase, Murota, & Hayama, 2020). Ticks are a major vector for animal diseases in Uganda. And, we have previously reported a high tick burden, comprising of several tick species on cattle under this study (Balinandi et al., 2020). Taken together, the high tick burden with high diversity of tick species parasitizing cattle provide avenues for likely transmission of several infectious agents, some of which may be novel.

Indeed, from our study, we report 4 novel viruses with unknown epidemiology in Uganda. Unfortunately, a major limitation of metagenomics is that, beyond detection, we are unable to determine the pathogenicity or medical-veterinary importance of the identified novel viruses. However, their presence in cattle blood indicates potential for spillover into the environment through vectors, or other biological and mechanical mechanisms. Of particular interest, was the identification of a pestivirus in our study, which we have tentatively named as Zikole virus. Pestiviruses are a group of reemerging viruses that have gained significant interest due to their wide geographical distribution, associated high socioeconomic losses when in livestock, a variety of syndromic manifestations in affected animals, failures in eradication programs, and a growing number of newly identified domestic and wildlife hosts (Blome, Beer, & Wernike, 2017; Postel, Smith, & Becher, 2021). Currently, they are known to infect a variety of artiodactyls, including swine and ruminants, in which they cause mild to severe disease (Schweizer & Peterhans, 2014). The Zikole virus identified in this study, had a close phylogenetic relationship to Bat pestivirus. While this may represent a mere common ancestry between these two viral species, recent studies have implicated bats as a major reservoir from which several known and unknown viral species have spilledover into other animal species (Wang & Anderson, 2019). From our study, the significance and impact of the identified Zikole virus on cattle, as well as human health remains unknown and a subject for further investigation. What is known, however, is that the common species of the genus *Pestivirus*, Bovine viral diarhea virus type 1, now re-classified as *Pestivirus A* (Smith et al., 2017) causes serious disease in cattle (Liebler-Tenorio, Kenklies, Greiser-Wilke, Makoschey, & Pohlenz, 2006).

As much as some of the viruses identified in this study are being reported for the first time in Uganda, they are known to be widely spread elsewhere in African bovines (Corman et al., 2015; Sibhat, Ayelet, Gebremedhin, Skjerve, & Asmare, 2018), where their clinical manifestations can range from absent overt disease to abortions, stillbirths and/or congenital malformations (Baechlein et al., 2015; Peperkamp et al., 2015). In this study, all virus-positive animals displayed normal rectal temperatures (Table 2), perhaps, indicating their tolerance to these viral infections (Ayres & Schneider, 2012). Nevertheless, a detailed characterization of these viruses is still crucial in the context of risk analysis, design of public health surveillance systems, and outbreak investigation and response. Of concern is the fact that many, if not all, of the identified viruses in the present study still lack effective vaccines or antiviral treatments. They are of unknown epidemiology, including their circulation in nature, range of potential natural reservoirs and immunological responses in relation to clinical infections, whether in man or in animals. Thus, the main limitation of this study emanated from its cross-sectional nature, which hindered us from obtaining detailed information regarding these epidemiological perspectives.

## Ethical Statement

The authors confirm that they read and adhrered to journal’s ethical policies as stated in the authors guidelines. The required ethical clearances to collect ticks and blood samples from cattle were obtained from the animal ethical committee and a research permit issued by the Uganda National Council of Science and Technology (UNCST).

## Data Availability Statement

Data supporting all components of this study is available on request from the corresponding author. Sequence data was submitted to GeneBank and can be accessed using accession numbers provided in the body of the text.

## CRediT Authorship Contribution Statement

**Stephen Balinandi**: investigation, methodology, visualization, writing - original draft **Juliette Hayer**: methodology, data curation, formal analysis, writing - review and editing **Harindranath Cholleti**: methodology, data curation, investigation, writing - review and editing **Michelle Wille**: methodology, data curation, formal analysis, visualization, writing - review and editing **Julius J. Lutwama**: investigation, supervision, writing - review and editing, **Maja Malmberg**: conceptualization, funding acquisition, investigation, methodology, data curation, formal analysis, resources, supervision, project administration, writing - review and editing, **Lawrence Mugisha**: conceptualization, investigation, supervision, project administarion, resources, methodology, visualisation, formal analysis, original draft, writing - review and editing, validation.

## Acknowledgements

We thank Dr. Bernard Ssebide from the Mountain Gorilla Veterinary and USAID/PREDICT projects in Uganda, for providing pan-PCR primers that were used in this study. In addition, sequencing was performed by the SNP&SEQ Technology Platform in Uppsala, Sweden. The facility is part of the National Genomics Infrastructure and Science for Life Laboratory. The SNP&SEQ Platform is also supported by the Swedish Research Council and the Knut and Alice Wallenberg Foundation. We also thank Conservation and Ecosystem Health Alliance (CEHA) tha coordinated the field sample collection activities together with livestock farmers that availed their animals for the study.

